# Neural dynamics become significantly faster and more nonlinear at the peak of information processing

**DOI:** 10.64898/2026.11.01.763548

**Authors:** Toktam Samiei, Sidd Lokray, Hafiz Fareed Ahmed, Erfan Nozari

## Abstract

The temporal scale at which neural populations preserve stimulus information is often treated as fixed, even though population activity changes rapidly during sensory processing. Whether the information-optimal scale changes within a single stimulus-evoked episode, and what determines its evolution, remains unclear. We developed Integration–Split, a decoding-based framework that estimates the time-varying temporal resolution of population spike trains. Across four visual paradigms and throughout the mouse visual hierarchy, the optimal scale contracted after stimulus onset and subsequently re-expanded. This pattern persisted across stimulus temporal frequencies and was largely absent in a hippocampal control. The optimal scale varied inversely with stimulus decodability, reaching its finest resolution when stimulus identity was most accurately decoded. Beyond this association, independently estimating the signal–noise autocorrelation balance across candidate window widths identified the window that maximized the autocorrelation-based signal-to-noise ratio, and this optimal width closely followed the Integration–Split trajectory. Consistent with these findings, stronger nonlinear predictive structure coincided with finer optimal temporal resolution, consistent with the idea that shorter integration windows are needed to preserve informative nonlinear population structure that would otherwise be attenuated by temporal averaging. To test the functional relevance of this organization, we trained a Transformer solely to decode stimulus identity, without providing the optimal temporal-scale pattern or Integration– Split outputs. Its learned attention exhibited a similar contraction-and-recovery profile, and disrupting heads with stronger correspondence produced larger decoding deficits. Together, these findings reveal a dynamically changing, information-optimal temporal scale in population coding, shaped by the evolving balance of signal and noise autocorrelation and by nonlinear population structure, and independently recapitulated in the learned attention of a Transformer trained without temporal-scale supervision.

## Introduction

Understanding how neural populations encode information requires identifying not only which activity patterns carry information, but also the temporal scale over which those patterns should be integrated. This scale determines how long recent activity remains relevant, which successive events are combined, and which rapid changes are preserved rather than smoothed away. Its functional importance is reflected in the cortical hierarchy of neural timescales: sensory regions exhibit shorter windows, whereas association and prefrontal areas exhibit progressively longer windows suited to integrating information over extended periods [1–3] Neural timescales also vary with consequential changes in brain state, including the induction and recovery of anesthesia [4], and atypical regional timescales have been associated with altered information processing and symptom severity in autism [5]. Thus, temporal scale is a fundamental feature of neural information processing with relevance to perception, cognition, consciousness, and disease.

Studies of neural timescale have typically estimated one value for a neuron, brain region, or experimental condition. Murray and colleagues, for example, assigned each cortical area one decay constant estimated over prestimulus foreperiods lasting approximately 0.5–1.5 s [2]. Other studies reported differences across attention conditions and trials [6], across minutes-long periods of behavioral engagement [7], and continuously during anesthesia transitions using 2-min sliding windows [4]. Together, these studies have characterized variation across neurons, regions, conditions, and relatively slow changes in brain state. It remains unknown, however, whether the relevant temporal scale also changes within the tens to hundreds of milliseconds over which a single sensory population response unfolds. Population codes can reorganize substantially within this brief interval [8–12]; consequently, a timescale estimated across the full response may conceal meaningful internal dynamics. If successive population states change rapidly, the duration over which activity can be integrated without mixing distinct representations may need to change with them. This raises two related questions: does the information-optimal temporal scale vary within a few hundred milliseconds, and which properties of the evolving population code determine that variation?

One candidate determinant is the relative temporal organization of signal and noise. Here, signal multivariate autocorrelation measures the persistence of stimulus-conditioned population patterns across temporal lags, whereas noise multivariate autocorrelation measures the temporal structure of trial-to-trial residual population activity. Their joint structure determines whether pooling activity improves or degrades population decoding [13–23]. Our theoretical work showed that the temporal scale optimizing stimulus information depends on the temporal balance between stimulus-related signal and trial-to-trial noise [24]. An evolving balance between signal and noise autocorrelation should therefore produce an evolving optimal temporal scale. In particular, when signal autocorrelation decays rapidly across temporal lags, longer integration increasingly mixes distinct stimulus-related population states, favoring finer temporal resolution.

The signal–noise autocorrelation balance provides one potential mechanism for dynamic temporal scaling, but population nonlinearity may provide another. Temporal and spatial averaging smooth population activity by attenuating nonlinear structure, making the observed dynamics progressively more linear [25–28]. Although such smoothing can improve robustness, it can also discard stimulus-specific information carried by nonlinear interactions, which can support high-dimensional representations and enhance separability [29]. The appropriate averaging scale should therefore adapt to the evolving degree of nonlinearity: finer resolution is needed when information-bearing nonlinear structure is strong, whereas broader integration may be beneficial when that structure is weak and averaging can improve reliability without erasing relevant detail. Together, signal–noise autocorrelation and nonlinear predictive structure provide candidate population-level mechanisms for explaining why the decoding-optimal scale should change during a response.

Testing this possibility requires an approach that estimates temporal scale locally from its effect on information, rather than imposing one integration window on the entire trial. It must allow neighboring portions of the same response to be represented at different resolutions while preventing changes in decoder dimensionality from determining the result. Large-scale Neuropixels recordings from the Allen Institute provide the population size, repeated stimulus presentations, and coverage of the mouse visual hierarchy needed for this analysis [30, 31].

Here, we developed *Integration–Split*, an optimization framework that adaptively integrates or separates neighboring portions of population spike trains according to held-out stimulus-decoding performance. We applied it to the dorsal lateral geniculate nucleus (LGd), primary visual cortex (V1), higher visual areas (HVAs), and the hippocampal formation (HPF) across Gabor patches, natural scenes, drifting gratings, and a visual change-detection task. To examine the previously uncharacterized within-response regime, we estimated the optimal temporal scale throughout a brief 250-ms response. The scale contracted rapidly after stimulus onset, reached its finest resolution near peak stimulus information, and then re-expanded. Thus, the information-preserving scale was reorganized not only across brain regions or behavioral states, but within a fraction of a second as the population code evolved. This trajectory varied inversely with stimulus decodability across visual regions but remained comparatively flat in HPF. The same organization persisted across drifting-grating frequencies, arguing against simple entrainment to stimulus tempo. Signal and noise multivariate autocorrelation explained why the scale contracted: near peak information, the autocorrelation-based signal-to-noise ratio was maximized by shorter windows, and its maximizing width closely tracked the Integration–Split solution. Nonlinear predictive advantage increased over the same interval and was likewise associated with finer temporal resolution. These results support a framework in which the optimal temporal scale is dynamically organized by the evolving signal–noise autocorrelation balance and nonlinear structure of the population response.

The temporal-scale estimates above were obtained by explicitly optimizing temporal partitions for stimulus decoding. To test whether similar temporal organization emerges under a different decoding architecture without temporal-scale supervision, we trained a Transformer solely to identify the stimulus from population spike trains. The network had no knowledge of the Integration–Split temporal-scale pattern; its temporal weighting was learned from the stimulus-decoding objective alone. We then asked whether the learned attention profiles resembled the Integration–Split trajectory and whether heads with stronger correspondence contributed more to decoding. Attention profiles exhibited a similar contraction-and-recovery pattern, and replacing the learned temporal weights of individual heads with uniform weights produced larger decoding deficits for heads that more closely matched the reference. These results provide convergent evidence for dynamic temporal organization beyond the original partitioning procedure and demonstrate the functional contribution of learned temporal weighting within the trained network.

## Results

### An optimization framework for estimating timescale dynamics in population spiking activity

In order to objectively estimate the— potentially dynamic—timescale of neural information processing on a millisecond-by-millisecond basis (Figure 1a), we developed an optimization framework consisting of two complementary components (Figure 1b, see Methods for details). The first is an end-to-end decoding module, established previously in [32], that quantifies the task-relevant information contained in population spiking activity by measuring how accurately a normative decoder identifies the presented visual stimulus (See Methods and [32]). Immediately *upstream* of the decoder, the *Integration–Split* module adaptively partitions the spike trains into variable-width temporal intervals (Figure 1c). This module iteratively merges neighboring temporal intervals or splits an existing interval and retains each operation according to whether the resulting representation lowers cross-validated decoding error (Figure 1d).

**Figure 1:**
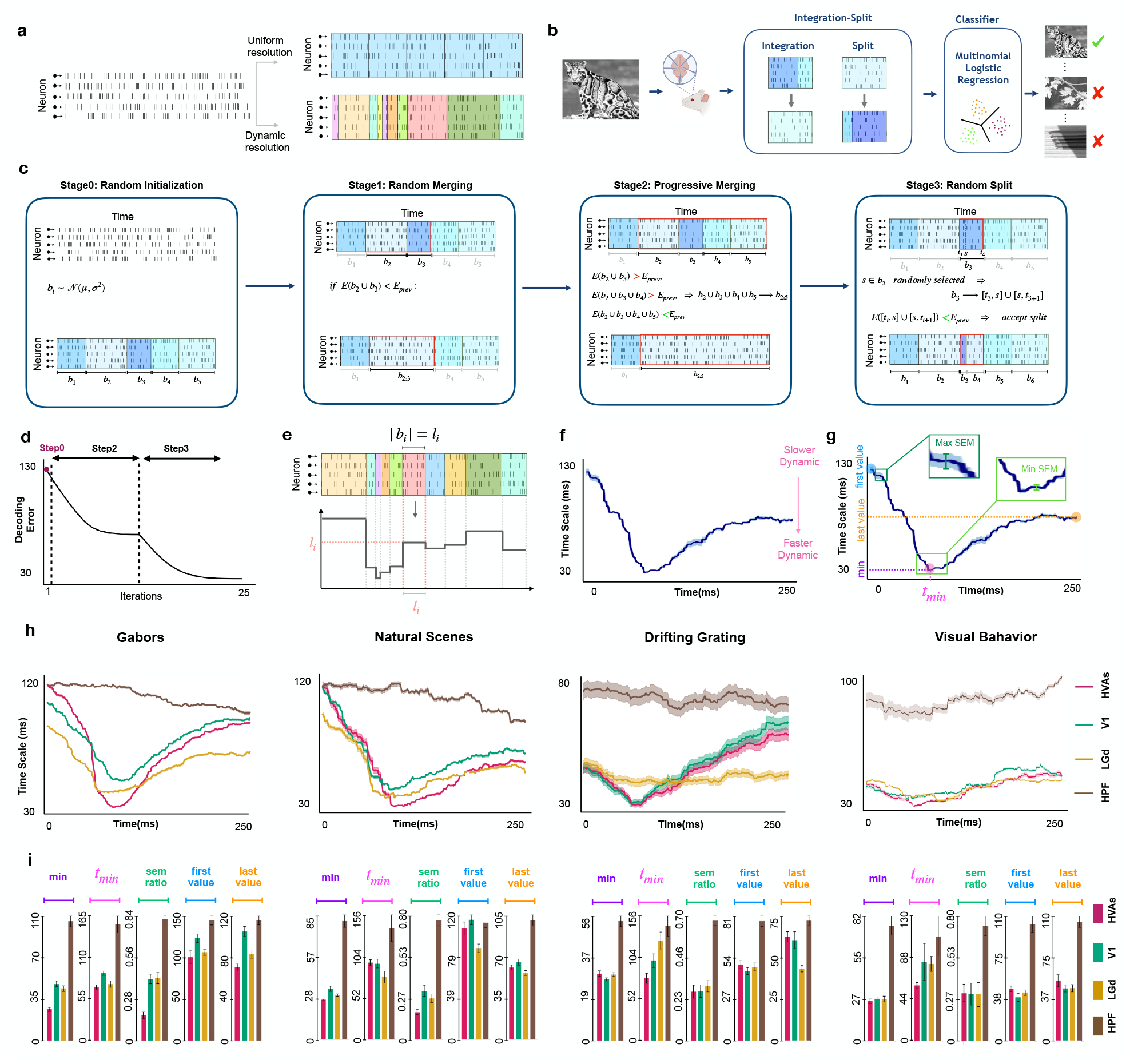
Integration–Split reveals a dynamic, decoding-optimal temporal scale. (a) Example population spike raster and two possible representations of the same activity: uniform fixed-width bins (*top*) or a time-varying partition of variable-width bins (*bottom*). (b) Analysis pipeline. Population spike trains evoked by a visual stimulus are temporally partitioned by Integration–Split and supplied to a multinomial logistic-regression decoder. Candidate integrations and splits are retained only when they reduce held-out classification error. (c) One representative step from each optimization stage is illustrated for a single spike-count matrix. *Stage 0*, initialization with bin widths sampled from *N*(*µ, σ*^2^). *Stage 1*, pairwise integration of adjacent bins. *Stage 2*, progressive directional integration: beginning with two adjacent bins, the candidate is expanded one neighboring bin at a time until the first integration that reduces decoding error is found, after which the search restarts from the updated partition. *Stage 3*, subdivision of a bin at a candidate split point *s*. Red outlines identify the bins being evaluated. (d) Held-out decoding error across optimization iterations for an example case. Error decreases as accepted integrations and splits accumulate and then stabilizes. (e) Conversion of the final partition into a continuous temporal-scale profile. For each bin *B*_*i*_ of width *l*_*i*_, the value *l*_*i*_ is assigned to every millisecond contained in that bin. (f) Mean ±SEM temporal-scale profile across 25 independent optimization runs for an example case. The estimated optimal scale contracts after stimulus onset and subsequently re-expands; smaller values indicate finer temporal resolution. (g) Summary measurements extracted from each profile: minimum scale, its time *t*_min_, initial and final scales, and the minimum and maximum SEM used to calculate the SEM ratio. (h) Mean ± SEM optimal temporal-scale profiles for Gabors, Natural Scenes, Drifting Gratings, and Visual Behavior, shown for HVAs (magenta), V1 (teal), LGd (gold), and HPF (brown). The three visual regions exhibit a contraction-and-recovery profile, whereas HPF shows a comparatively coarse and weakly modulated trajectory. (i) Minimum scale, *t*_min_, SEM ratio, initial scale, and final scale summarized by region for each stimulus paradigm (mean *±* SEM).

The resulting optimal partition was subsequently converted into an optimal timescale trajectory by assigning to each time point the duration of the interval to which it belongs (Figure 1e). This entire Integration–Split procedure was then repeated 25 times using independent random seeds, and the resulting trajectories were averaged to obtain a robust timescale estimate for each subject, brain region, and task condition (Figure 1f). Together, these steps estimate the temporal organization of information representation directly from the population response, adaptively determining where activity can be combined over longer intervals and where finer temporal distinctions must be preserved.

### The neural timescale is highly dynamic during early visual processing

We applied the above optimization pipeline on large-scale single-unit NeuroPixels spiking activity from four brain regions (3 visual, 1 control) across 16 mice (Gabors, Natural Scenes, and Drifting Gratings) and 10 mice (Visual Behavior), spanning four visual tasks (Figure 1h, see also Methods). Across all visual regions—the dorsal lateral geniculate complex (LGd), primary visual cortex (V1), and higher visual areas (HVAs)—and task conditions, the neural timescale changes rapidly and substantially within the brief 250-ms trial duration. In particular, the neural timescale initially decreases (becomes faster) post stimulus onset, before it again increases (becomes slower) until the end of the trial period. This pattern was robust across static (Gabors, Natural Scenes, and Visual Behavior) and dynamic (Drifting Grating) stimuli, suggesting that the observed dynamics of neural timescale reflect rapid changes in the temporal organization of the population response during sensory processing. We further repeated the same analysis pipeline on population activity in the hippocampal formation (HPF), which serves as a control region given its lack of low-level visual information encoding targeted by the decoder. As expected, the resulting timescale trajectories did not show the same dynamics as visual regions, and were instead relatively flat throughout the trial duration. These results thus demonstrate the presence of rapid fluctuations not only in the neural population response itself, but also in the timescale of such response, during early visual processing.

### Visual regions share a common trajectory of dynamic neural timescale

Although the full Integration–Split procedure was applied to each region independently, the resulting timescale trajectories recovered from visual regions followed a largely similar trajectory (Figure 1h). Across all four stimulus paradigms, the visual regions transitioned from relatively coarse temporal scales to a pronounced minimum before recovering toward coarser scales. Although the absolute temporal scales and the precise timing of their minima varied across regions and stimulus paradigms, this contraction-and-recovery profile was consistently expressed throughout the visual pathway.

To quantify the degree of similarity among timescale trajectories, we extracted and compared their initial, final, and minimum values as well as the time at which their minimum occurs (*t*_min_) and the ratio of their smallest to largest variability (s.e.m. ratio) (Figure 1g). The results, shown in Figure 1i, demonstrate that all visual regions reach their fastest timescale during a narrow window approximately 60– 100 ms after stimulus onset, and that this minimum was generally bracketed by larger initial and final temporal scales. The subject-to-subject variability in timescale trajectory also tended to be lowest near *t*_min_, resulting in relatively low minimumto-maximum s.e.m. ratios (∼0.3) across independent subjects and tasks.

As noted earlier, the timescale trajectories of HPF as a non-visual-encoding control region remained considerably coarser and lacked the pronounced contraction-and-recovery profile observed in the visual regions (Figure 1h). Its minimum, initial, and final temporal scales were consistently higher, while its minimum-to-maximum SEM ratio remained closer to one (Figure 1i). This higher ratio indicates that the SEM varied relatively little across the trial, without the localized reduction near *t*_min_ observed in the visual regions. These pronounced differences between HPF and visual regions ensure that the dynamic trajectories observed in the latter are specific, and reflect underlying dynamics of evolving stimulusrelated activity throughout the thalamocortical visual processing hierarchy.

### Trajectories of neural timescale align with neural information content

Given the reproducible V-shaped profile of neural timescale trajectories observed above and the known timing of information representation in visual cortex [33], we hypothesized that the former’s initial rapid decrease and subsequent slow increase may coincide with the initial arrival of visual information and the subsequent homeostatic normalization, respectively. To test this hypothesis, we analyzed whether the faster timescales were associated with higher stimulus decodability, and vice versa. To obtain the dynamic trajectory of stimulus decodability, we applied a short sliding-window analysis, as shown in Figure 2a. A separate decoder was trained and evaluated at each successive (sliding) 25-ms window, and its F1 score was tracked throughout the trial period (Figure 2b). As expected, stimulus decodability varied substantially during the trial duration, even though the visual stimuli remained present throughout the trial duration in all cases (Figure 2c). Across all task conditions, stimulus information increased rapidly following a short delay (∼50 ms) after stimulus onset, reached a pronounced and relatively narrow peak, and subsequently declined.

**Figure 2:**
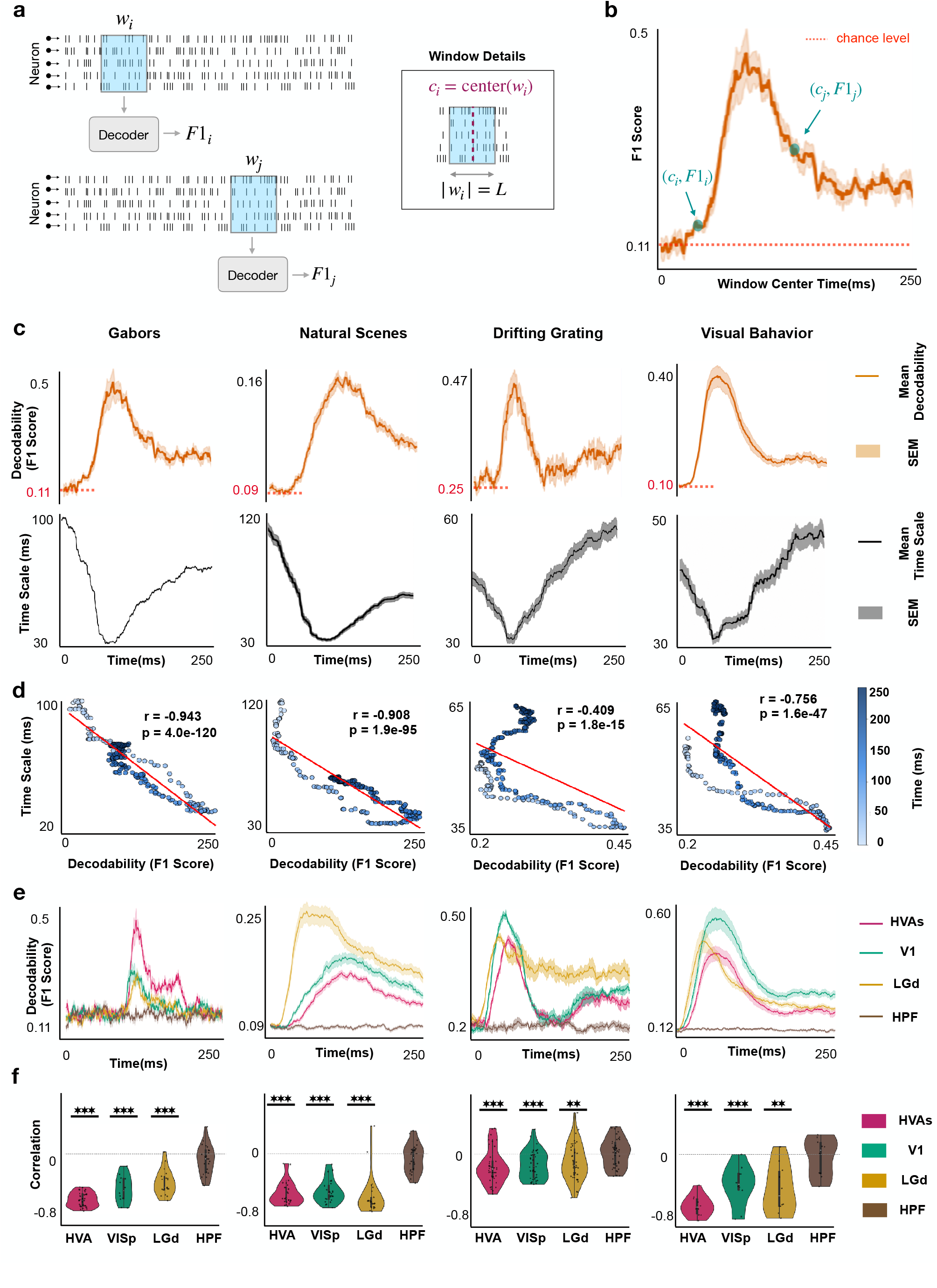
Stimulus decodability varies inversely with the optimal temporal scale. (a) Sliding-window decoding procedure. The neuron-by-time spike-train segment within a fixed 25-ms window *w*_*i*_, centered at *c*_*i*_, is supplied to a multinomial logistic-regression decoder to obtain the macro-averaged F1 score *F* 1_*i*_. The window is advanced in 1-ms steps to generate a time-resolved decodability profile. (b) Example F1-score profile for one classification case. Decodability rises from chance level (red dotted line), reaches a transient peak, and then declines; two example window centers are marked. (c) Mean ±SEM decodability (*top*, orange) and Integration–Split optimal temporal scale (*bottom*, black) in HVAs for each stimulus paradigm. Higher decodability generally accompanies a finer optimal temporal scale. (d) Optimal temporal scale plotted against decodability across time for the corresponding HVA profiles. Points are colored by time, red lines show least-squares fits, and Pearson correlations are *r* = *−*0.943 for Gabors, *−*0.908 for Natural Scenes, *−*0.409 for Drifting Gratings, and *−*0.756 for Visual Behavior. (e) Mean ±SEM decodability profiles separated by region: HVAs (magenta), V1 (teal), LGd (gold), and HPF (brown). Visual regions show stimulus-evoked increases in decodability, whereas HPF remains near chance. (f) Distributions of the per-case Pearson correlation between decodability and optimal temporal scale for each region and paradigm. Points denote individual pseudo-session/classification cases. Asterisks indicate a correlation distribution different from zero by a two-sided one-sample *t*-test (^*\**^*p <* 0.05, ^*\*\**^*p <* 0.01, ^*\*\*\**^*p <* 0.001; no multiple-comparison correction).

Consistent with our hypothesis, the time course of stimulus decodability exhibited a strong negative correlation with neural timescale. Across all task conditions, the initial rapid increase in decodability aligned with the rapid decrease in neural timescale, the two reached their extreme values nearly simultaneously, and relaxed back towards baseline after-wards (Figure 2c). As shown in Figure 2d for HVAs, the trajectories of decodability and neural timescale were strongly negatively correlated across all tasks (|*r*| *>* 0.4, *p <* 10^*−*15^, Student’s t-test), particularly in Gabors and natural scenes (|*r* |*>* 0.9). These findings confirm that the neural timescales identified by the Integration–Split procedure are closely aligned with the evolving amount of stimulus-related information present in the neural population.

Both the dynamic profiles of stimulus decodability and their inverse relationship with neural timescale were consistent across visual brain regions, and absent in the hippocampus (Figure 2e,f). Across LGd, V1, and HVAs decodability increased shortly after stimulus onset and subsequently declined, although the magnitude and duration of the information-bearing response differed across regions and stimulus conditions. In contrast, HPF remained close to chance-level decoding and showed little systematic temporal modulation. Consistent with these profiles, the correlation between decodability and neural timescale was strongly negative across visual regions but near zero in HPF.

Given the reliance of the estimated neural timescale and decodability trajectories on the same underlying end-to-end decoding architecture, we also tested this relationship using information-theoretic entropy instead of decodability. Unlike decodability, entropy is calculated without relying on any decoders or even regard for class labels, and instead quantifies information by the extent of uncertainty that each realization resolves [15, 19, 34]. Nevertheless, as shown in Supplementary Figure S1, the time course of entropy closely follows that of decodability, and hence similarly correlates negatively and significantly with neural timescale in visual regions but not the hippocampus.

While the neural timescale closely aligned with the time course of stimulus decodability, their alignment is correlational and does not imply a causal relationship in either direction. In other words, the alignment between the two does not explain *why* preserving stimulus information needed a finer temporal resolution during the peak of decodability and vice versa. This observation motivated us to examine the statistical structure of information representation across the trial duration, as described next.

### Dynamics of signal and noise multivariate autocorrelation explain the trajectories of neural timescale

Motivated by our recent theoretical work demonstrating the causal effect of signal and noise autocorrelation structure on the optimal *static* timescale of neural representation [24], we asked whether the same can explain our observed *dynamic* trajectories of neural timescale. In [24] we showed the presence of a mechanistic link between the relative temporal structure of stimulus-related information (‘signal’, Figure 3a) and trial-to-trial variability (‘noise’, Figure 3b), and the optimal scale of temporal integration. Specifically, whether integrating neighboring population responses improves or impairs stimulus discriminability depends on the *relative* decay rates of signal and noise autocorrelation functions, with larger integration windows (slower timescales) corresponding to slower signal autocorrelation decay compared to noise. Their relative temporal structure therefore determines whether adjacent responses should be integrated or preserved separately. Motivated by this prediction, we asked whether the lag-resolved multivariate autocorrelation functions of signal and noise evolve throughout the trial and whether their joint dynamics account for the temporal scale selected by the Integration–Split framework.

**Figure 3:**
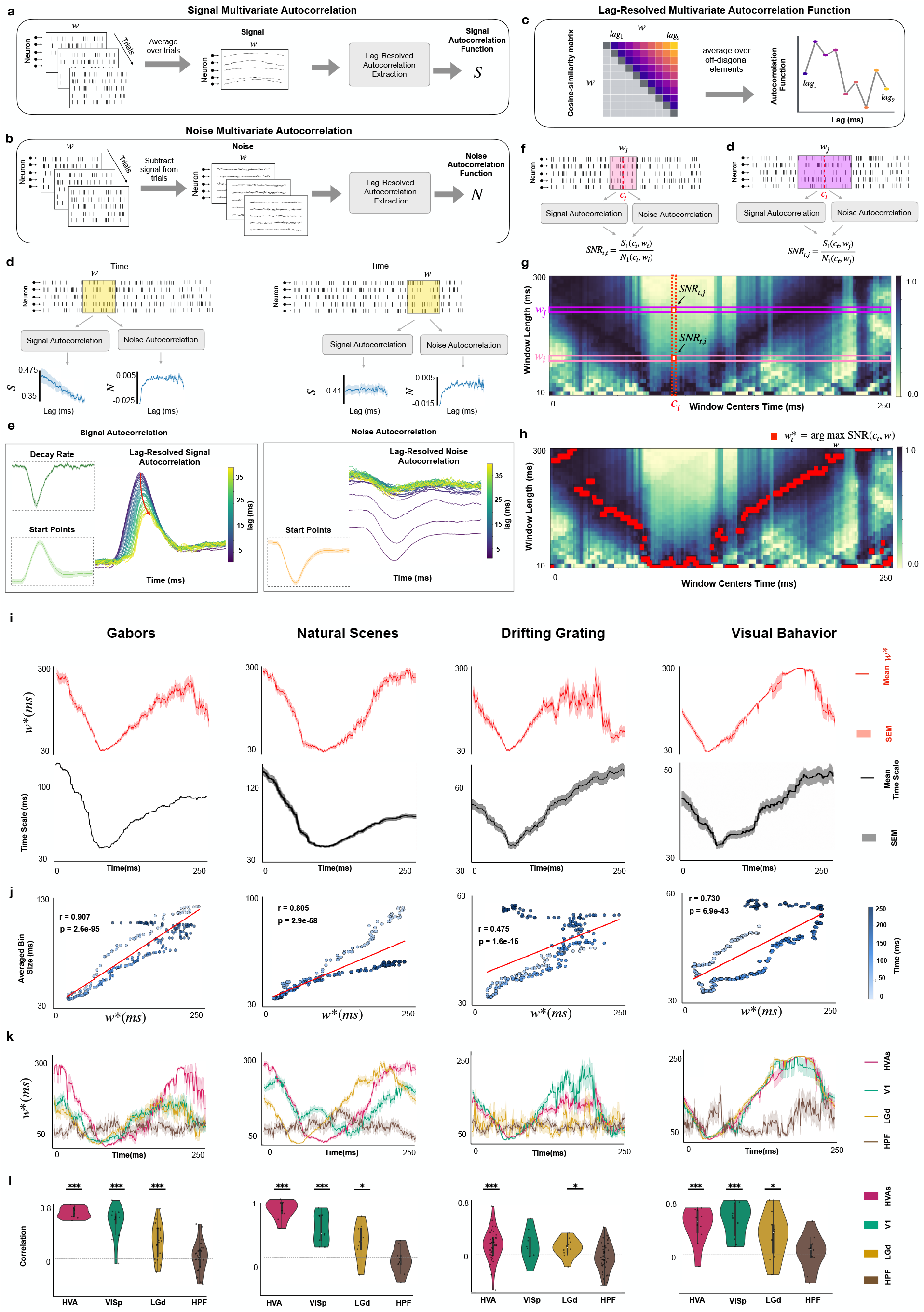
The evolving balance of signal and noise multivariate autocorrelation tracks the optimal temporal scale. (a) Signal multivariate autocorrelation. Population responses are averaged across trials within each stimulus class, and a lag-resolved autocorrelation function is extracted from the resulting class-mean population states. (b) Noise multivariate autocorrelation. The class mean is subtracted from each trial, and a lag-resolved autocorrelation function is extracted from the residual population activity. (c) Lag-resolved multivariate autocorrelation extraction within a window of width *w*. Pairwise similarities between population states form a matrix (cosine similarity for signal and Pearson correlation for noise); averaging entries at each temporal lag yields the corresponding lag-resolved autocorrelation function. The first off-diagonal defines the shortest-lag values *S*_1_ and *N*_1_. (d,f) Two windows of different widths, *w*_*j*_ and *w*_*i*_, centered at the same time *c*_*t*_. Their signal and noise autocorrelation functions yield the autocorrelation-based signal-to-noise ratios SNR_*t,j*_ = *S*_1_(*c*_*t*_, *w*_*j*_ )*/N*_1_(*c*_*t*_, *w*_*j*_ ) and SNR_*t,i*_ = *S*_1_(*c*_*t*_, *w*_*i*_)*/N*_1_(*c*_*t*_, *w*_*i*_). (e) Lag-resolved signal (*left* ) and noise (*right* ) multivariate autocorrelation functions across the trial. Curves are colored by temporal lag; insets show the shortest-lag starting values and the decay rate of signal autocorrelation. Near peak information, signal autocorrelation is generally higher and noise autocorrelation generally lower across the examined lags; signal autocorrelation also exhibits a steeper decline with increasing lag. (g) Autocorrelation-based signal-to-noise ratio as a function of window center and width for an example case. The two example windows and their ratio values from (d,f) are indicated. (h) At each center time, the red trace marks the window width that maximizes the autocorrelation-based signal-to-noise ratio, 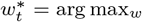 SNR(*c*_*t*_, *w*). (i) Mean ± SEM 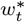 (*top*, red) and Integration–Split optimal temporal scale (*bottom*, black) for each stimulus paradigm. (j) Integration–Split temporal scale plotted against 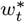 across time. Pearson correlations are *r* = 0.907 for Gabors, 0.805 for Natural Scenes, 0.475 for Drifting Gratings, and 0.730 for Visual Behavior. (k) Mean ± SEM 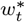 profiles separated by region: HVAs, V1, LGd, and HPF. (l) Distributions of the per-case Pearson correlation between 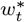 and the Integration–Split scale. Points denote individual pseudo-session/classification cases. Asterisks indicate a correlation distribution different from zero by a two-sided one-sample *t*-test (^*\**^*p <* 0.05, ^*\*\**^*p <* 0.01, ^*\*\*\**^*p <* 0.001; no multiple-comparison correction).

To test for an analogous dynamic relationship in real data, we estimated population-level signal and noise multivariate autocorrelation functions within sliding windows across the trial (Figure 3a–d). For signal, cosine similarity quantified the alignment of class-mean population states at pairs of time points; for noise, correlation quantified the alignment of the corresponding residual population activity. Averaging entries with the same temporal separation produced each lag-resolved autocorrelation function (Figure 3c). Both functions varied substantially between periods close to and far from the peak of information (Figure 3d,e). Near the information peak (∼70 ms), signal autocorrelation was strongest at short lags and decreased markedly with increasing lag, whereas later in the trial (∼150 ms) it remained comparatively flat across lags. Noise autocorrelation, by contrast, was negative at short lags and gradually increased with lag; its shortest-lag value became more negative near the information peak. Thus, peak information was associated not with a uniform shift in signal or noise autocorrelation, but with a sharper lag-dependent organization of both components: stronger signal autocorrelation and more negative noise autocorrelation between neighboring time points, together with a steeper decay of signal auto-correlation over longer lags.

We next tested whether the temporal evolution of signal and noise autocorrelations can explain the observed trajectories of neural timescale [24]. Let *S*_1_(*c*_*t*_, *w*) and *N*_1_(*c*_*t*_, *w*) denote the shortest-lag signal and noise autocorrelation values for a window of length *w* centered at *c*_*t*_, respectively. We defined the autocorrelation-based signal-to-noise ratio

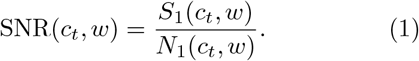

as illustrated in Figure 3d,f. We calculated SNR(*c*_*t*_, *w*) by continuously varying *c*_*t*_ throughout the trial and discretely varying *w* from 10 to 300 ms in 10-ms increments (Figure 3g). At each fixed *c*_*t*_, we then identified the width 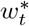 that maximized this ratio (Figure 3h). Near peak information, SNR(*c*_*t*_, *w*) was maximized at substantially shorter widths than later in the trial (*c*_*t*_ ∼ 200 ms), indicating that the evolving signal and noise autocorrelation structure could account for the dynamic neural-timescale trajectory. As the response progressed, the maximum shifted toward longer windows, indicating an increasing advantage of integrating activity over broader temporal intervals.

As hypothesized, the dynamics of 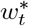 strongly and positively predicted those of the neural timescale (Figure 3i). Across all task conditions, both trajectories decreased toward finer timescales near the peak of information and subsequently increased to-ward longer integration windows. Their temporal correlations were positive and significant in all four paradigms (Figure 3j; *r* = 0.475–0.907, *p <* 10^*−*15^, Student’s t-test). The same relationship was reproduced across visually informative brain regions and subjects, whereas HPF exhibited comparatively flat 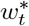 trajectories that lacked significant correlations with neural timescale (Figure 3k,l).

Together, these findings provide a mechanistic account of dynamic neural timescale through the relative lag-resolved strength of signal and noise multi-variate autocorrelation. The remaining question is why the autocorrelation-based signal-to-noise ratio itself favors a finer scale near peak information; we address this next.

### Dynamic nonlinearity explains the trajectories of neural timescale

Motivated by prior work demonstrating a strong deteriorating effect of spatiotemporal integration on the nonlinearity of neural dynamics [25–28], we hypothesize that dynamic nonlinearity should *increase* near the peak of information, therefore giving rise to “more to be lost” through (excessive) integration. In other words, we hypothesize that strongly nonlinear dynamics require finer temporal integration (faster timescale) in order to preserve their stimulus-representing nonlinearities, whereas approximately linear dynamics can be integrated over longer intervals—and benefit from cancellation of weakly correlated noise—without losing nonlinear structure.

To test this hypothesis, we compared linear and nonlinear predictive models of population response over a sliding window of constant length (Figure 4a– c). The linear model described future population response (future state) through a *linear superposition* of current population response (state) and stimulus-encoding components (input), whereas the nonlinear model allowed for predicting the former via potentially nonlinear combinations of the latter two. We then quantified the extent of nonlinearity present in each window via

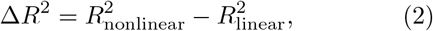

where larger values of Δ*R*^2^ indicate a greater improvement in prediction when nonlinear interactions are included.

**Figure 4:**
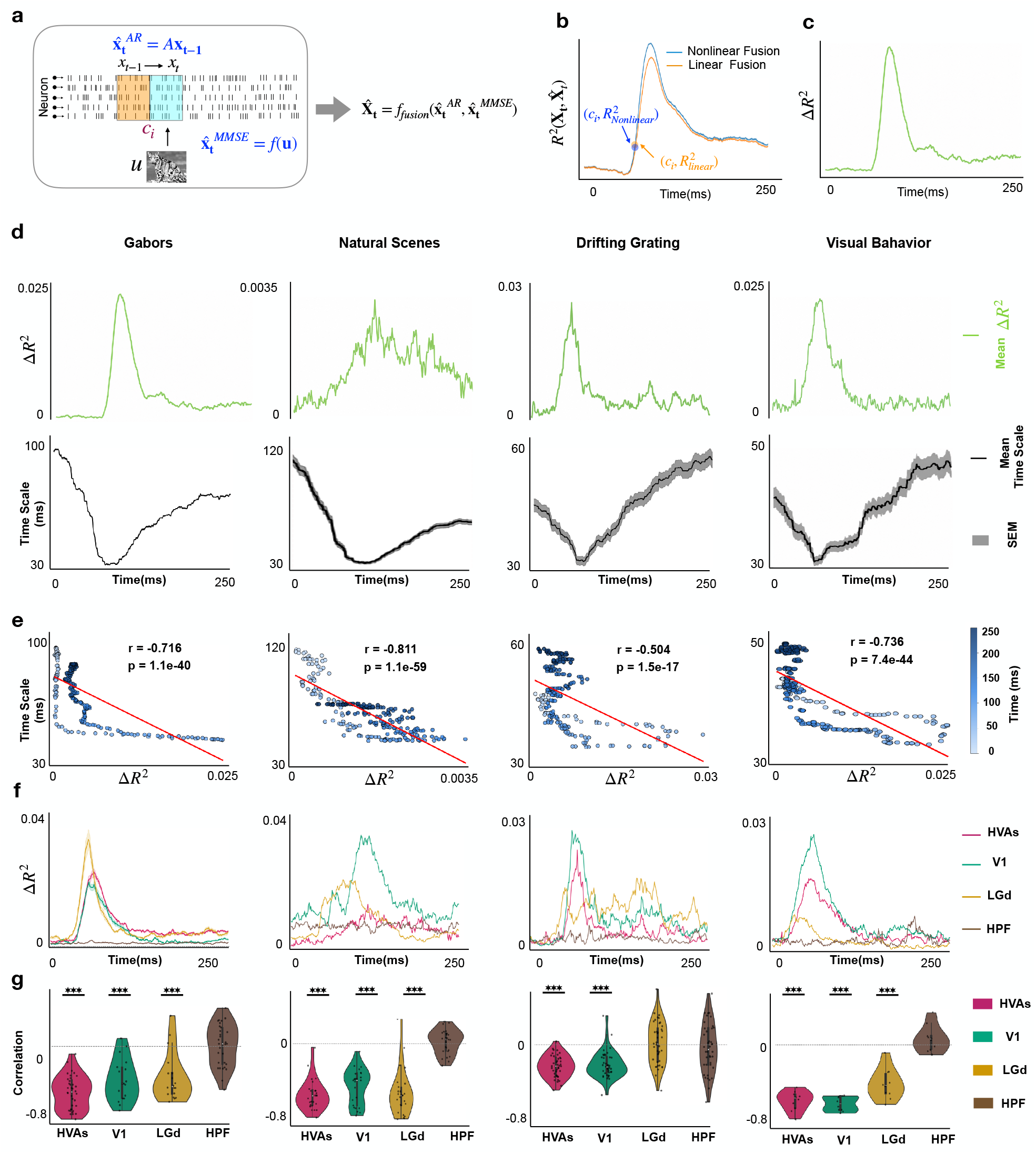
Nonlinear predictive advantage is greatest when the optimal temporal scale is finest. (a) Linear–nonlinear prediction framework. Adjacent bins define the recent population state, *X*_*t−*1_, and target response, *X*_*t*_, while *U* denotes the stimulus features. A linear autoregressive history predictor, 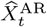, and a nonlinear stimulus-feature predictor, 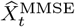, provide complementary component predictions. Linear fusion combines these components by per-neuron linear regression, whereas nonlinear fusion additionally models gated nonlinear residual interactions (Methods). (b) Example held-out variance explained, *R*^2^, for the linear-fusion (orange) and nonlinear-fusion (blue) models across the trial. (c) Nonlinear predictive advantage for the same example, defined as 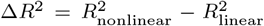 . Positive values indicate an improvement in held-out prediction from nonlinear fusion relative to linear fusion. (d) Mean *±* SEM Δ*R* (*top*, green) and Integration–Split optimal temporal scale (*bottom*, black) for each stimulus paradigm. Nonlinear advantage increases near peak information as the optimal scale contracts. (e) Optimal temporal scale plotted against Δ*R*^2^ across time. Pearson correlations are *r* = *−*0.716 for Gabors, *−*0.811 for Natural Scenes, *−*0.504 for Drifting Gratings, and *−*0.736 for Visual Behavior. (f) Mean ± SEM Δ*R*^2^ profiles separated by region: HVAs, V1, LGd, and HPF. (g) Distributions of the per-case Pearson correlation between Δ*R*^2^ and optimal temporal scale. Points denote individual pseudo-session/classification cases. Asterisks indicate a correlation distribution different from zero by a two-sided one-sample *t*-test (^*\**^*p <* 0.05, ^*\*\**^*p <* 0.01, ^*\*\*\**^*p <* 0.001; no multiple-comparison correction).

Across all task conditions, the amount of dynamic nonlinearity varied substantially throughout the trial, with Δ*R*^2^ increasing as stimulus decodability rose and reaching its maximum near the peak of information (Figure 4d).

As with decodability (Figure 2e), the neural timescale was negatively correlated with Δ*R*^2^ across time in every task condition (Figure 4e; | *r*| *>* 0.5, *p <* 10^*−*15^, Student’s t-test), most strongly in Gabors, Natural Scenes, and Visual Behavior (| *r* | *>* 0.7). This relationship was consistent across visual brain regions and absent in the hippocampus (Figure 4f,g). Across LGd, V1, and HVAs, Δ*R*^2^ increased shortly after stimulus onset and subsequently declined, tracking the same time course as decodability and the inverse of neural timescale, whereas HPF showed negligible Δ*R*^2^ throughout (Figure 4f). Consistent with these profiles, the correlation between Δ*R*^2^ and neural timescale was strongly negative across visual regions but near zero in HPF (Figure 4g).

Together, these findings indicate that the dynamic trajectory of neural timescale is governed not only by the amount of decodable information and the balance between signal and noise, but more fundamentally by the degree of nonlinearity in the population dynamics; information-rich periods with stronger nonlinear contributions require finer temporal resolutions, while periods with weaker nonlinear structure permit broader temporal integration.

### A Transformer independently trained to decode stimulus identity learns functionally relevant temporal organization

Transformers [35] can learn to selectively weight informative components of their inputs through self-attention, providing an alternative approach for investigating the temporal organization of neural population activity. Motivated by this capability, we asked whether the dynamic temporal-scale trajectory identified by Integration–Split could also emerge from a Transformer trained solely to decode stimulus identity, without any explicit temporal-scale supervision. To test this, we trained a Transformer to identify Gabor stimuli from HVA population spike trains. The architecture consisted of two encoder layers, each containing four attention heads, with spike events represented by their neuron identities and occurrence times (Figure 5a). Unlike causal attention, the bidirectional self-attention used here allowed each spike event to attend to both earlier and later events. Moreover, the multi-head architecture provided an opportunity to examine different temporal-weighting patterns learned simultaneously by separate attention heads and, through selective head ablation, assess their functional contribution to decoding. We therefore investigated whether the learned attention patterns resembled the Integration–Split trajectory and whether heads showing stronger correspondence were more important for stimulus decoding.

**Figure 5:**
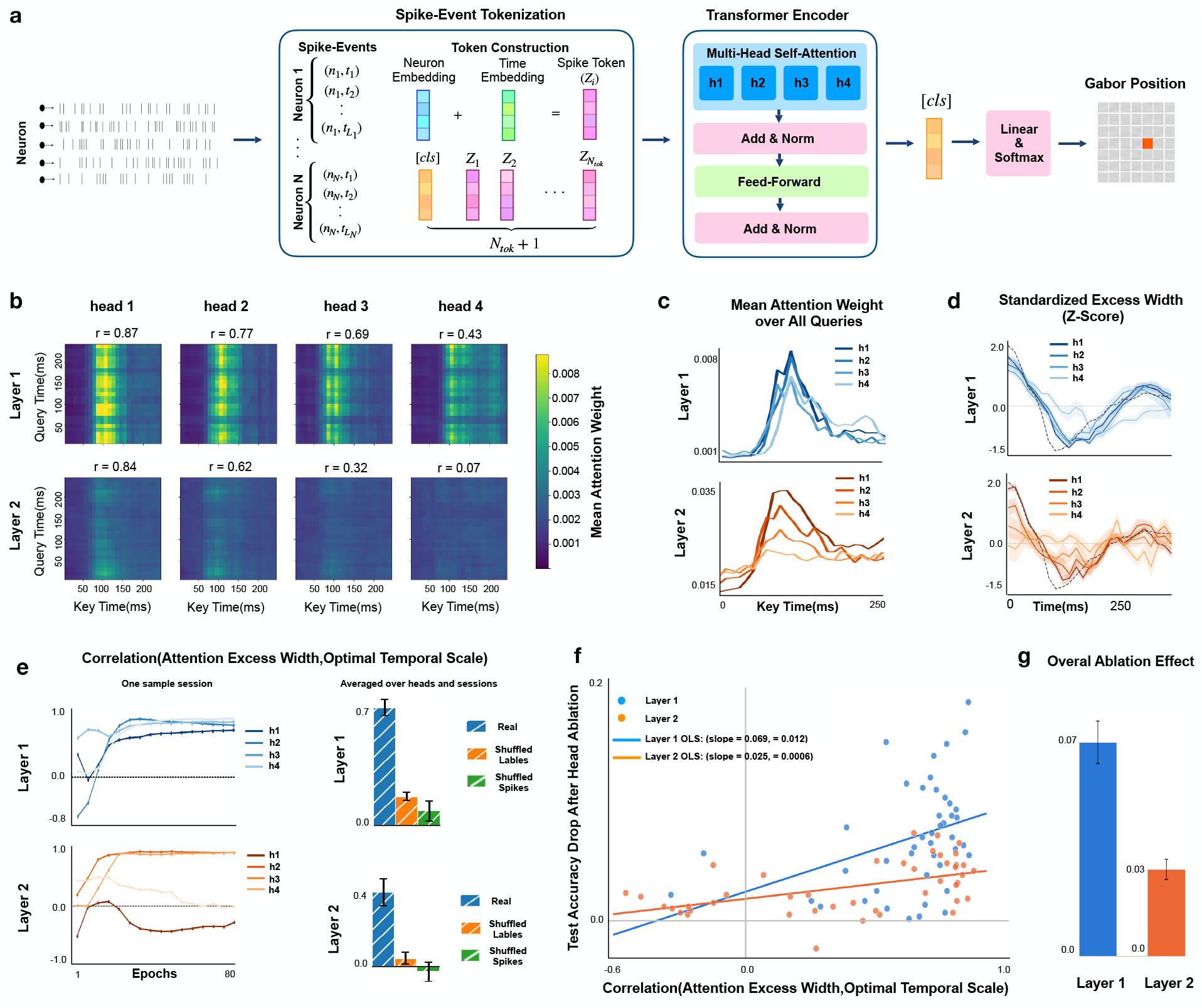
An independently trained Transformer learns dynamic temporal organization that supports stimulus decoding. **(a)** Neuron-identity and time embeddings represent each spike event. A two-layer, four-head bidirectional Transformer uses a classification token ([CLS]) to predict Gabor position on the 9 × 9 grid. Models were trained independently on HVA recordings without a temporal-scale objective or access to Integration–Split outputs. **(b)** Query-by-key attention maps averaged across sessions. Within each session and layer, heads were ranked by the Pearson correlation between their attention excess-width profile and the Integration–Split optimal temporal-scale reference, then averaged across sessions by rank. Head 1 denotes the most strongly correlated head. Reported *r* values are the mean session-level correlations for each rank. Vertical bands indicate preferential weighting of particular stimulus-relative times across queries. **(c)** Attention weights averaged over query times for the corresponding ranked heads. **(d)** Standardized excess-width profiles for the corresponding ranked heads, alongside the Integration–Split reference (black dashed curve). Excess width is the attention-weighted mean absolute query– key time separation minus its uniform-attention expectation. First-layer profiles resemble the contraction-and-recovery pattern of the reference more consistently than second-layer profiles. **(e)** Correlation between excess width and the reference during training in an example session (left), and averages across heads and sessions for real data, shuffled-label controls, and shuffled-spike controls (right). The reference was digitized from the example Integration–Split trajectory in Figure 1f. **(f)** Single-head ablation replaces learned attention weights with uniform weights over valid keys while preserving the value pathway. Each point represents one head in one session. Greater correspondence with the reference is associated with a larger decrease in held-out decoding accuracy (pooled Spearman *ρ* = 0.491, *p* = 1.2 × 10^*−*6^; 88 head–session observations across 11 sessions and both layers). Lines show layer-specific ordinary least-squares fits. **(g)** Mean decrease in held-out accuracy following individual-head ablation, summarized by layer. Layers 1 and 2 denote the first and second encoder layers, respectively.

Attention maps obtained from 11 independently trained session-specific models revealed prominent vertical bands, particularly in the first encoder layer, indicating that specific key times received attention from queries distributed throughout the target interval (Figure 5b). Averaging attention weights over queries further revealed preferential weighting of the informative poststimulus interval, consistent with the temporal profile of stimulus decodability identified by sliding-window analysis (Figures 5c and 2). While these attention maps indicated when the network preferentially attended, they did not directly characterize the temporal extent over which activity was integrated. To quantify this property, we defined the attention excess width as the attention-weighted mean absolute temporal separation between queries and keys, after subtracting the expected separation under uniform attention (see Methods).

The resulting standardized excess-width profiles exhibited a contraction-and-recovery trajectory resembling the optimal temporal scale identified by Integration–Split, particularly in the first encoder layer (Figure 5d). However, the strength of this correspondence varied across attention heads and layers. To characterize this variability, we ranked heads within each session and layer according to the Pearson correlation between their excess-width profiles and the Integration–Split reference trajectory, and subsequently averaged their profiles across sessions by rank. The mean session-level correlations for the four ranked heads were *r* = 0.87, 0.77, 0.69, and 0.43 in the first layer, compared with *r* = 0.84, 0.62, 0.32, and 0.07 in the second layer (Figure 5b). Thus, although both layers contained heads whose temporal organization closely resembled the Integration–Split trajectory, this correspondence was more consistently expressed across heads in the first layer.

We next examined how this temporal organization developed during training and whether it depended on learning stimulus-related information. In an example session, first-layer heads progressively converged toward positive correlations with the Integration–Split reference, whereas second-layer heads exhibited more heterogeneous training trajectories (Figure 5e, left). Averaged across heads and sessions, the correspondence was substantially stronger for models trained on real data than for those trained with shuffled stimulus labels or shuffled spikes (Figure 5e, right). Furthermore, randomly initialized, untrained networks exhibited excess-width variations that were five-to tenfold smaller in magnitude and lacked a coherent temporal trajectory across sessions. Together, these control analyses indicate that the observed temporal organization emerges through learning from stimulus-related neural activity rather than from the network architecture alone.

We further investigated whether the excess-width trajectories could be explained by simpler properties of attention or stimulus information. First, we examined whether the temporal width at each query was determined by its distance from the most-attended key time. This relationship was generally weak (mean per-curve *r* = 0.14; first layer: *r* = 0.22; second layer: *r* ≈ 0.05), with only 8% of curves exhibiting correlations above 0.7. Second, we compared the excess-width profiles with local stimulus decodability estimated by sliding-window analysis. Excess width was negatively correlated with decodability in the first layer (mean *r* = −0.42), whereas the second layer showed little average linear association (mean *r* ≈ 0.03). These findings suggest that the learned temporal-width trajectories cannot be fully explained by either distance from a preferentially attended time or their linear relationship with local stimulus decodability.

Although the similarity between the attention excess-width profiles and the Integration–Split trajectory suggests that the Transformer recovered a comparable temporal organization, this correspondence alone does not establish its functional importance for decoding. We therefore performed a single-head ablation analysis [36] to test whether heads showing stronger correspondence contributed more to the model’s decoding performance. Specifically, we replaced the learned attention weights of one head at a time with uniform weights over valid keys, while preserving its value projections and all other model parameters. We then quantified the resulting decrease in held-out stimulus-decoding accuracy. As hypothesized, heads whose excess-width profiles were more strongly correlated with the Integration–Split reference produced larger accuracy losses when ablated (Figure 5f; pooled Spearman *ρ* = 0.491, *p* = 1.2 × 10^*−*6^, *n* = 88 head–session observations). This association was significant in both the first (*ρ* = 0.439, *p* = 0.0029, *n* = 44) and second (*ρ* = 0.496, *p* = 6.3 × 10^*−*4^, *n* = 44) encoder layers, despite the weaker average relationship between excess width and local decodability in the second layer. Furthermore, the mean decoding-accuracy loss following individual-head ablation was greater in the first layer than in the second (Figure 5g).

Together, these findings demonstrate that a Transformer trained exclusively to decode stimulus identity can develop temporal attention patterns resembling the dynamic optimal temporal scale identified by Integration–Split, without explicit supervision of this organization. Importantly, the ablation analysis further shows that heads exhibiting stronger correspondence contribute more to stimulus decoding, supporting the functional relevance of the learned temporal weighting. Although the excess-width measure is distinct from the optimal integration window estimated by Integration–Split, and the ablation perturbs the entire attention-weighting pattern rather than selectively removing its contraction-and-recovery component, these results provide convergent evidence that similar temporal organization can emerge from an independently trained decoding architecture.

## Discussion

### Summary

We identified rapid changes in the decoding-optimal temporal scale of stimulus-evoked neural population activity during early visual processing. Across four visual paradigms, the optimal scale contracted shortly after stimulus onset and subsequently recovered, with its minimum occurring near peak stimulus decodability. The scale was negatively associated with both decodability and nonlinear predictive advantage, and positively associated with the window width maximizing an autocorrelation-based signal–noise measure. These relationships were consistently observed in visual regions but were weak or absent in the hippocampal control. A Transformer trained independently for stimulus classification developed attention-width profiles with similar temporal organization; heads whose profiles more closely matched the Integration–Split trajectory also tended to contribute more to decoding. Together, these findings identify dynamic temporal resolution as a feature of stimulus-related population coding and motivate hypotheses about the population statistics underlying its evolution.

### Potential implications among the observed dynamic trends

Our analyses reveal coordinated changes in decoding-optimal temporal scale, stimulus decodability, signal and noise autocorrelation, and nonlinear predictive advantage. These associations do not establish a causal sequence, but they motivate a possible account of why shorter integration windows are favored near peak information. In particular, temporal averaging can attenuate nonlinear dynamical structure [25–28] If some stimulus-related distinctions depend on nonlinear population interactions, integrating activity over excessively long windows could obscure those distinctions, making them less accessible to the decoder. In such a regime, the loss of potentially informative structure could out-weigh the benefits of averaging trial-to-trial variability, favoring finer temporal resolution. Conversely, when the relevant structure changes more slowly or nonlinear predictive contributions are weaker, longer integration windows may improve reliability without sacrificing as much discriminative information. Our observation that nonlinear predictive advantage increases as the optimal temporal scale contracts is consistent with this possibility, although it does not demonstrate that the measured nonlinear component itself carries stimulus-specific information.

The autocorrelation analyses suggest two related but distinct aspects of the information-rich response. First, as illustrated in Figure 3e, signal autocorrelation generally increased and noise autocorrelation generally decreased near peak information across the examined temporal lags. This opposing shift is consistent with a more favorable relationship between stimulus-related population structure and trial-to-trial fluctuations at a fixed temporal scale, potentially contributing to higher stimulus decodability. Second, the dependence of autocorrelation on temporal lag is relevant to the optimal integration window. Near peak information, signal autocorrelation declined more steeply as the lag increased, indicating that population states separated by longer intervals were less similar than neighboring states. Broad temporal integration could therefore combine increasingly distinct stimulus-related patterns. Consistent with this interpretation, the width 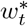 maximizing the autocorrelation-based measure became shorter and closely tracked the Integration–Split trajectory. The all-lag changes in signal and noise help interpret the decodability peak, whereas their lag-dependent organization helps interpret why a finer integration scale is favored.

One possibility is that the transient increase in non-linear predictive structure contributes to this reorganization of signal and noise autocorrelation, with consequences for both decodability and optimal temporal resolution (Figure 6). Alternatively, all three changes could arise in parallel from the evolving stimulus-evoked population response. Our observations do not distinguish these explanations, and testing the proposed causal relationships will require targeted manipulations or models that isolate the relevant population interactions.

**Figure 6:**
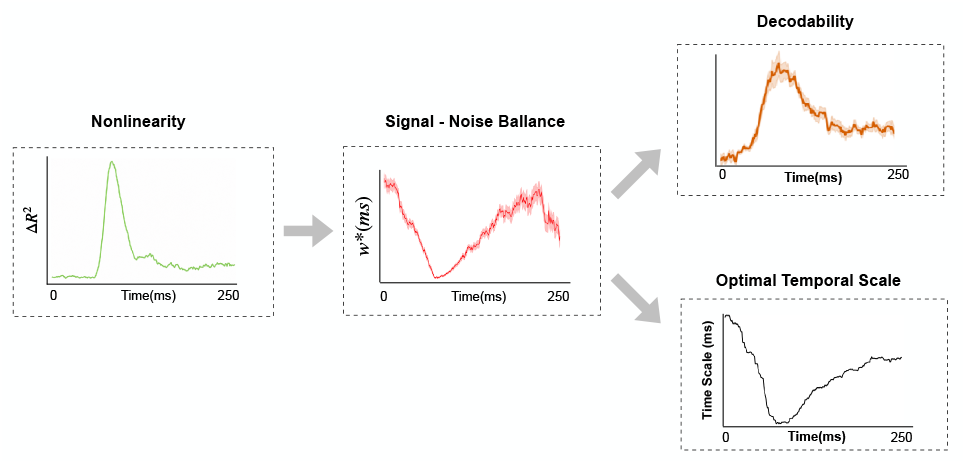
Proposed mechanistic organization linking nonlinearity, signal–noise autocorrelation, decodability, and optimal temporal scale. The hypothesized pathway links stronger nonlinear predictive structure near peak information to higher signal and lower noise autocorrelation across the examined lags, together with a steeper decline in signal autocorrelation as temporal lag increases. The lag-dependent organization is proposed to favor a shorter autocorrelation-based signal-to-noise-maximizing window, 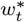 (*center* ). The overall change in signal relative to noise could support greater stimulus separability and decodability (*top right* ), whereas the shorter 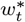 is associated with a finer Integration–Split optimal temporal scale (*bottom right* ). Example profiles over the same 250-ms epoch illustrate the coordinated temporal relationships among the four quantities.

### Response entropy tracks temporal scale but not stimulus-specific information

Response entropy provided a label-independent measure of population-response variability. Like decodability, entropy increased near the time at which the optimal temporal scale contracted and subsequently declined (Supplementary Figure S1). Thus, the relationship between response variability and temporal scale can be observed without using stimulus labels or training a decoder for the entropy calculation.

However, entropy and stimulus decodability were not interchangeable. After the information peak, entropy fell below its prestimulus level even though decoding performance remained above chance. Entropy describes the uncertainty of the observed response distribution, irrespective of which stimulus was presented; decodability describes the ability to distinguish stimulus classes from those responses [15]. Distinct but reliable stimulus-evoked states can therefore support decoding even when response entropy is low, whereas high entropy can arise from stimulus-independent variability. We accordingly use decodability as the primary measure of stimulus-specific information and interpret entropy as a complementary, label-independent description of response variability.

#### Linear self-dynamics, nonlinear stimulus encoding, and their evolving superposition

Our predictive model combines two complementary sources of information about the population response (Figure 4). The linear autoregressive predictor, *Ŷ*^*AR*^, describes how recent activity predicts the subsequent population state, whereas the stimulus-based predictor, *Ŷ*^*MMSE*^, estimates the response from visual input features. This separation is motivated by evidence that macroscopic population dynamics can often be approximated by linear models [25, 28], even when stimulus-to-response mappings are nonlinear [37].

The difference Δ*R*^2^ measures how much additional held-out response variance the nonlinear fusion model explains beyond linear fusion of these same predictors. It is therefore an operational measure of nonlinear predictive advantage, not a direct estimate of intrinsic circuit nonlinearity or stimulus-specific information. Its increase near peak decodability indicates that nonlinear fusion is especially useful during that part of the response. Together with the simultaneous contraction of the decoding-optimal temporal scale, this finding supports the hypothesis that the usefulness of temporal integration depends in part on the evolving nonlinear predictive structure of the population response; it does not establish that these changes causally determine one another.

### Temporal-frequency independence of the optimal temporal scale

One alternative explanation is that the changing optimal temporal scale simply reflects entrainment to the stimulus’s temporal structure. Drifting gratings provided a test of this possibility because they were presented at five temporal frequencies (1, 2, 4, 8, and 15 Hz). If the optimal temporal scale directly tracked stimulus periodicity, faster gratings might consistently favor shorter integration windows. Instead, the visual regions displayed a similar contraction-and-recovery trajectory across frequencies, without a consistent ordering of optimal scales by frequency (Supplementary Figure S2). The absence of a systematic frequency ordering argues against a simple correspondence between grating frequency and decoding-optimal temporal scale. It does not exclude more complex effects of stimulus dynamics, but it is consistent with a stimulus-onset-locked reorganization of the information represented in the population response.

### Controlling for decoder dimensionality

A potential confound in optimizing temporal partitions is that splitting or merging bins can change the number of features provided to a decoder. We addressed this by using fixed-size populations of 50 neurons for the Integration–Split analyses and by keeping the decoder input length constant across candidate partitions (Methods). Specifically, each bin’s mean neural activity was repeated at every elementary time point within the bin before the resulting array was flat-tened for decoding. The partition therefore changed the temporal resolution of the representation without changing its nominal feature count. This control does not make the representations equivalent in effective dimensionality: temporal averaging necessarily changes their covariance and fine-scale structure. Those changes are part of the effect of the representation being tested, rather than a consequence of simply providing a different number of input features.

### Scope and limitations

The temporal scale estimated here is defined by the preservation of stimulus identity under a particular decoder and is not an independent measurement of a circuit’s intrinsic autocorrelation time constant or biological integration window. Its absolute value may depend on the decoder, stimulus set, trial count, population size, and binning constraints. The Integration–Split algorithm also provides an approximate rather than globally optimal partition: its greedy search explores a finite set of merges and splits and depends partly on its initialization. Repeating the procedure with independent seeds and data splits yielded consistent contraction-and-recovery profiles, supporting the robustness of the temporal pattern while leaving individual bin boundaries and absolute scale estimates approximate.

The reported relationships are observational. In particular, the autocorrelation-based measure and Δ*R*^2^ characterize specific population statistics and predictive models; neither alone establishes a causal mechanism for the change in decoding-optimal scale. Correlations calculated between closely spaced time points should also be interpreted primarily as descriptive measures of trajectory alignment, because those observations are temporally dependent. Moreover, pseudo-sessions sampled from the same recording share biological and experimental context, and the time-resolved profiles are estimated across repeated presentations rather than demonstrating trial-by-trial adjustment of a biological integration window. Finally, single-head attention ablation tests the contribution of a head’s learned weighting as a whole; it does not selectively isolate the contraction- and-recovery component of that weighting.

### Conclusion

Our findings reveal a rapidly changing, decoding-optimal temporal scale during early visual processing. Temporal resolution becomes finer near peak stimulus information and subsequently broadens, while signal and noise autocorrelation and nonlinear predictive advantage undergo coordinated changes. A separately trained Transformer recovers related temporal attention patterns, providing a complementary perspective on this organization. Together, these results suggest that the temporal resolution most useful for population decoding depends on the momentary structure of the neural response. Identifying the circuit mechanisms underlying these associations, and testing whether similar dynamics occur in other sensory, cognitive, and motor systems, remain important directions for future work.

## Methods

### Data

Neural recordings were obtained from two publicly available Allen Institute Brain Observatory Neuropixels datasets. The Visual Coding dataset contains large-scale extracellular recordings from cortical and subcortical regions in awake, head-fixed mice during passive presentation of a standardized visual-stimulus battery [30]. We analyzed the Gabor stimuli from the functional-connectivity stimulus set, natural scenes, and drifting gratings. These passive-viewing analyses comprised recordings from 16 mice. The companion Visual Behavior Neuropixels dataset contains recordings acquired while mice performed a visual image change-detection task [31]; 10 sessions met the fixed-population criterion described below. Spiking data and stimulus metadata were accessed through the AllenSDK `EcephysProjectCache` interface. No new animal experiments were performed for this study.

### Brain-region definitions

Units were assigned to anatomical groups from their Allen Common Coordinate Framework structure acronyms. Analyses were performed separately for higher visual areas (HVAs), primary visual cortex (V1), the dorsal lateral geniculate nucleus (LGd), and the hippocampal formation (HPF); the constituent structures are listed in Table 1. HPF was included as a non-visual control because it did not exhibit reliable stimulus-evoked decoding in these data. The same regional definitions were used for all four stimulus paradigms.

**Table 1:** Brain regions analyzed in this study and their constituent Allen CCF structures.

| **Region** | **Constituent structures** |
|---|---|
| HVAs | VISal, VISam, VISl, VISli, VISmma, VISpm, VISrl |
| V1 | VISp |
| LGd | LGd |
| HPF | CA1, CA2, CA3, DG, SUB |

### Spike preprocessing and fixed-size populations

Spike preprocessing and trial reconstruction followed previously described procedures [38]; all parameters used in the present analysis are specified below. Sorted single-unit spike times were obtained directly from each dataset and discretized into 1-ms bins. To make decoder dimensionality comparable across sessions, regions, and stimulus conditions, all analyses used populations of *N* = 50 neurons. Within each session and region, well-isolated units were randomly sampled without replacement in non-overlapping blocks of 50 using a session-specific random seed. Each block was treated as an independent pseudo-session, allowing recordings with more than 50 eligible units to contribute multiple non-overlapping populations without discarding surplus units. Session–region combinations with fewer than 50 units were excluded.

For each train/test split, the training-trial mean of every neuron was subtracted from both training and test data; test trials therefore did not contribute to centering. No temporal filtering, smoothing, or trial shuffling was applied unless explicitly stated for a particular analysis.

### Trial construction

Each analyzed trial comprised three consecutive 250-ms stimulus presentations from the presentation-ordered stimulus table: the presentation preceding the event of interest, the event of interest, and the following presentation. Concatenating these epochs produced a 750-ms spike-count trace, with the analyzed stimulus occupying the middle interval, [250, 500) ms. This convention supplied temporal context around the 250-ms epoch reported in the main text and was applied to all four paradigms. For presentations at a session boundary, a randomly selected presentation from the same stimulus class was substituted for the missing neighbor.

### Organization of classification subtasks

Each classification case contained a non-overlapping subset of stimulus identities. Gabor classes were defined by patch position on the 9 × 9 spatial grid and divided into non-overlapping 9-class subtasks. For natural scenes, each image identity defined one class. Images were divided into 10 non-overlapping subtasks of 11 identities whose frame indices were spaced by 10; every identity had 50 repeated presentations, of which 40 were used for training and 10 for testing. For drifting gratings, the eight motion directions were divided into two four-class subtasks, with each pair of opposite directions corresponding to the same grating orientation assigned to different subtasks. This division was performed separately within each temporal-frequency condition (1, 2, 4, 8, and 15 Hz), yielding 10 drifting-grating classification problems in total. For Visual Behavior, the eight natural-scene identities in the change-detection task formed a single 8-class problem and were not subdivided further.

### Integration–Split optimization of temporal resolution

The Integration–Split algorithm estimates the temporal partition that best preserves stimulus identity for a population decoder (Figure 1b,c). A candidate partition of a trial of duration *T* was defined by bin edges

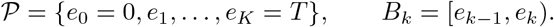

The partition *P* was shared across all trials and neurons. For each trial *q* and neuron *n*, activity was averaged within each bin *B*_*k*_, and the bin-specific mean was repeated at every millisecond *τ* ∈ *B*_*k*_:

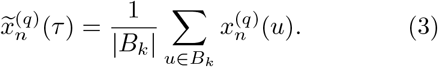

For each candidate partition *P*, the transformed activity of each trial was flattened into a decoder input vector in ℝ*NT* . Thus, merging or splitting bins changed only the temporal organization, while input dimensionality remained fixed.

#### Decoder and objective

For every candidate partition, an *l*_2_-penalized multinomial logistic-regression classifier was fit to the training trials using the `lbfgs` solver and warm-started from the preceding candidate. The regularization parameter was fixed at *C* = 0.1 for every region and task, selected from a candidate grid by decoding performance. The optimization objective was held-out classification error,

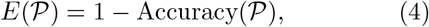

evaluated on the task-specific set of held-out trials. A proposed integration or split was retained only when it strictly decreased *E*(*P* ).

#### Initialization

The initial partition was generated by drawing successive bin widths from *N*(25 ms, 4^2^ ms^2^), rounding to the nearest millisecond, and enforcing a minimum width of 1 ms until the entire trial was covered.

#### Random pairwise integration

Adjacent bins were visited in random order and proposed for pairwise integration. An integration was accepted only when it reduced held-out decoding error. This stage continued for at most 10 full passes and terminated early when two consecutive passes left the partition unchanged.

#### Progressive directional integration

Starting from the pairwise-integration solution, the algorithm swept through the partition and tested progressively broader integrations, beginning with two adjacent bins and extending across as many as five additional neighboring bins. The first integration that reduced held-out decoding error was accepted, after which the search restarted from the updated partition. Left-to-right and right-to-left sweeps were both performed, and their order was alternated across independent optimization runs to avoid favoring either direction. This stage also continued for at most 10 passes or until convergence.

#### Splitting

Every interval remaining after integration was tested for subdivision. Candidate split locations were drawn from a Beta(2, 2) distribution over the interior of the interval, favoring locations near its center. The number of tested candidates increased with interval width, from 2 candidates for intervals no longer than 10 ms to 50 candidates for intervals longer than 100 ms. Intervals were visited in random order for at most 50 rounds, with each round comprising one complete pass through all intervals in the current partition. A split was retained only when it decreased decoding error. After five consecutive rounds without an accepted split, the algorithm performed a brief greedy reintegration over groups of at most three neighboring intervals before resuming the split search. Optimization ended when two consecutive complete passes produced no change.

#### Optimization ensemble and temporal-scale profile

The complete procedure was repeated 25 times for every region, pseudo-session, and classification subtask, using an independent train/test split and random seed on each repetition. For a final partition *P*, the time-resolved scale was defined as

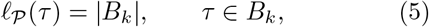

so that the width of a bin was repeated across every millisecond contained in that bin (Figure 1e). Smaller values indicate finer temporal resolution, whereas larger values indicate broader temporal integration. The 25 profiles were averaged to obtain the mean ± SEM profile for each case. The minimum scale, its time *t*_min_, the initial and final scales, and the ratio of the SEM at the minimum to the maximum SEM were calculated within the middle 250-ms epoch and then summarized across cases.

### Time-resolved stimulus decodability

Stimulus decodability was measured independently of the Integration–Split optimization using a fixed 25-ms sliding window (Figure 2a,b). At each window center, the neuron-by-time spike-train segment within the window was flattened directly to form one feature vector per trial. Features were standardized using the training data, and an *l*_2_-penalized multinomial logistic-regression classifier (`lbfgs` solver) was fit on a stratified 80/20 train/test split. Performance was quantified by the macro-averaged F1 score. The window was advanced in 1-ms steps across the trial, yielding a decodability profile. Profiles were averaged over 25 independent train/test partitions and are reported as mean ± SEM.

### Response entropy

As a label-independent measure of response diversity, we estimated the Shannon entropy of each neuron’s spike count within the same 25-ms sliding window. For each neuron *n* and window center *t*, spikes were summed within the window separately on every trial, yielding one spike-count value *r* per trial. The relative frequencies of the unique spike-count values across trials defined the empirical distribution *p*_*n*_(*r*| *t*) over *r* ∈ {0, 1, 2,*…*}. The entropy was

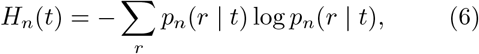

and population response entropy was the mean across the fixed population,

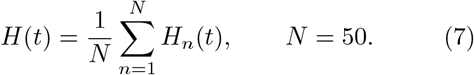

No finite-sample bias correction was applied.

### Signal and noise multivariate autocorrelation analysis

Signal and noise multivariate autocorrelation functions were estimated separately to determine how stimulus-related population structure and trial-to-trial residual activity changed across temporal lags (Figure 3a–c). Let **x**_*c,q*_(*u*) ∈ ℝ^*N*^ denote the population response at time *u* on trial *q* of stimulus class *c*. The stimulus-conditioned mean and residual responses were

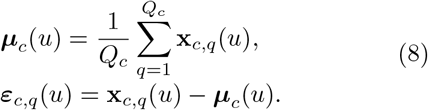

where *Q*_*c*_ is the number of trials in class *c*.

For two times *u* and *v* within a window of width *w* centered at *c*_*t*_, signal multivariate autocorrelation was quantified as the cosine similarity between class-mean population vectors,

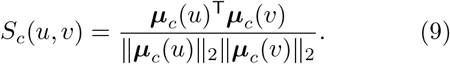

After pooling residuals across neurons and trials within each class, noise multivariate autocorrelation was quantified as their Pearson correlation, equivalent to cosine similarity because the residuals were zero-mean:

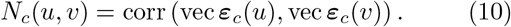

For each component, values with the same lag Δ = |*u* − *v*| were averaged along the corresponding matrix diagonal and then across stimulus classes. This produced the lag-resolved multivariate autocorrelation functions *S*(*c*_*t*_, *w*; Δ) and *N* (*c*_*t*_, *w*; Δ) for Δ = 1, …, *w* − 1.

The shortest-lag values, *S*_1_(*c*_*t*_, *w*) = *S*(*c*_*t*_, *w*; 1) and *N*_1_(*c*_*t*_, *w*) = *N* (*c*_*t*_, *w*; 1), were entered into the autocorrelation-based signal-to-noise ratio in Equation (1). The calculation was repeated for *w* ∈ {10, 20, …, 300} ms at every analyzed center time, producing the surface SNR(*c*_*t*_, *w*) shown in Figure 3g.

The window width that maximized this ratio was

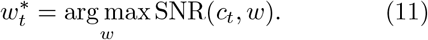

### Linear and nonlinear models of population dynamics

To quantify nonlinear predictive structure, we modeled the target population response, *X*_*t*_, from two complementary sources: the recent population state, *X*_*t−*1_, and the visual stimulus features, *U* (Figure 4a–c). At each evaluated target time step *t*, spike counts were summed within two adjacent, non-overlapping windows, each 25 ms in duration. The earlier window defined *X*_*t−*1_ ∈ ℝ^*N*^, and the later window defined *X*_*t*_ ∈ ℝ^*N*^ . The index *n* ∈ {1, …, *N*} denotes neuron.

#### Linear history predictor

For each neuron *n*, a univariate autoregressive model predicted the target response from that neuron’s activity in the preceding bin:

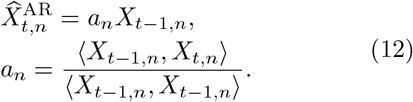

Both *X*_*t−*1_ and *X*_*t*_ were mean-centered using the training-set means, and the coefficient was estimated by ordinary least squares on the training trials with no intercept.

#### Stimulus-feature construction

For Gabor stimuli, the stimulus vector *U* contained orientation and spatial position (*x* and *y*). For drifting gratings, *U* contained orientation and temporal frequency. Natural scenes do not have an equivalent set of continuous experimental parameters; scene identity alone provides only a categorical label and therefore does not define a meaningful distance between images for the kernel regression used below. We instead derived a low-dimensional feature representation directly from the images. Each image was flattened into a pixel vector, and every pixel dimension was standardized across the natural-scene ensemble. A full principal-component analysis (PCA) was first fit to this standardized image matrix. The first five principal components, which together explained 28.27% of the total variance, were retained. PCA was then refit with five components, and the resulting component scores were standardized across images. For each natural-scene class *c*, these scores defined a feature vector *U*_*c*_ ∈ ℝ^5^ that was shared by all repeated trials belonging to that class. Across *C* natural-scene classes, the vectors formed a feature matrix *U*∈ ℝ*C×*^5^.

#### Nonlinear stimulus predictor

Each stimulus class *c* was represented by its corresponding feature vector *U*_*c*_ ∈ℝ^*d*^, with *d* = 3 for Gabor patches, *d* = 2 for drifting gratings, and *d* = 5 for natural scenes. The target response for a test stimulus *U* ^*\**^ was estimated from the training trials using Nadaraya–Watson regression with a Gaussian kernel:

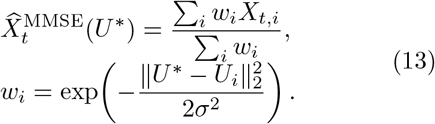

The bandwidth was fixed at *σ* = 0.1 × range(*U* ).

#### Linear and nonlinear fusion

An unconstrained per-neuron linear regression combined the history predictor, 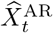, and stimulus predictor, 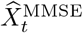, into the linear-fusion prediction, 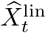:

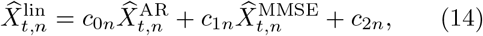

fit by ordinary least squares. A gated model then predicted the residual left by the linear-fusion model using neuron-specific saturating gates:

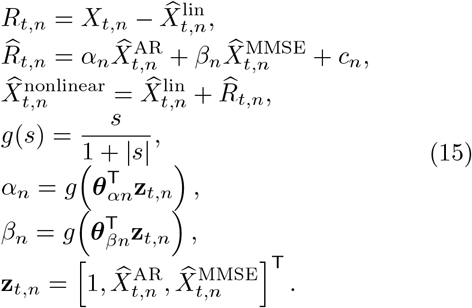

The neuron-specific fusion and gate parameters were estimated by L-BFGS-B minimization of the *l*_2_-regularized mean-squared residual error. Regularization strength was selected independently at each time offset by held-out performance from the grid *λ* ∈ {0, 10^*−*8^, …, 10^9^}.

Predictive performance was evaluated on the task-specific set of held-out trials using

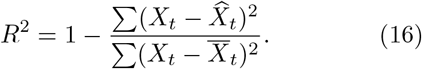

The linear and complete nonlinear models yielded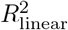 and 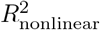, respectively. Their difference, Δ*R*^2^ (Equation (2)), measures the predictive advantage of nonlinear fusion. The analysis was repeated across time offsets and 25 independent train/test partitions, and the resulting profiles were averaged as mean ± SEM.

### Attention-based Transformer model of population spiking activity

A Transformer decoder [35] was trained to classify Gabor stimulus identity from HVA population spike trains (Figure 5a). The training objective contained only stimulus-classification loss; the Integration–Split temporal-scale reference was used exclusively for subsequent comparisons and head ranking. Models were trained independently for 13 sessions, of which 11 completed the attention-extraction and ablation analyses.

#### Population activity and spike-event tokens

All well-isolated HVA units were retained within each session, without the fixed 50-neuron subsampling used for Integration–Split. The input comprised the 250-ms target presentation and 250 ms of context on each side, giving a 750-ms sequence. The class label referred only to the target presentation. Let 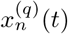 denote the spike count of neuron *n* in the 1-ms bin at time *t* on trial *q*. Each occupied neuron–time bin generated one event token, so counts greater than one were represented by a single event. For event *i* with neuron identity *n*_*i*_ and time *t*_*i*_, the token embedding was

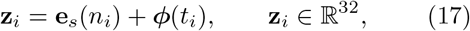

where **e**_*s*_ is the neuron-identity embedding learned independently for session *s*, and ***ϕ*** is the continuous sinusoidal time embedding shared across neurons and sessions. Times were measured relative to target-stimulus onset. A learned [CLS] token was prepended to the event sequence to provide a trial-level readout [39].

#### Encoder architecture and classification

The token sequence passed through two Transformer encoder layers, each with four attention heads, embedding dimension 32, feed-forward hidden dimension 64, and dropout 0.1 (Figure 5a). For layer *l* and head *h*, learned projections of the layer input provided query, key, and value matrices *Q*^*lh*^, *K*^*lh*^, and *V* ^*lh*^. Scaled dot-product attention was computed as

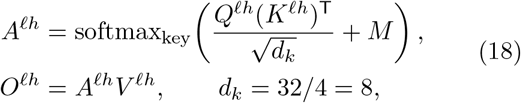

where *M* excludes masked keys and the softmax normalizes each query over its valid keys. Attention was bidirectional, without a causal restriction to earlier times. The four head outputs were concatenated and projected back to dimension 32. As shown in the architecture schematic, the attention and position-wise feed-forward sublayers were each followed by a residual addition and layer normalization [35]. The final [CLS] representation **c**^(*q*)^ was mapped to class probabilities by

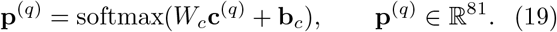

The output classes corresponded to all positions on the 9 × 9 Gabor grid, with class index 9 row + column for zero-based row and column indices. Thus, each model solved one 81-class problem, whereas Integration–Split used non-overlapping 9-class subtasks.

#### Training and evaluation

The 3645 trials per session (81 positions × 45 repeats) were divided into stratified training, validation, and test sets in proportions 70/15/15. Parameters were optimized using the mean training-set cross-entropy,

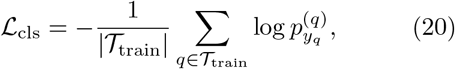

where *y*_*q*_ is the target stimulus class. Adam [40] used a learning rate of 3 × 10^*−*4^, weight decay 10^*−*4^, and batch size 16. Training ran for at most 200 epochs, with early stopping after 20 epochs without improvement in validation loss. No temporal-scale target or attention-profile matching term was included. Held-out classification accuracy was compared with a logistic-regression baseline using total spike counts and with chance performance of 1*/*81.

#### Attention maps and temporal width

For each session, layer, and head, attention weights on held-out trials were grouped into query–key time bins of 10 ms and averaged across trials (Figure 5b,c). Only spike-event tokens contributed to these maps; [CLS] and padding were excluded. Temporal profiles were evaluated for query times in the central 250-ms target interval, with neighboring presentations providing key context. Let *B*_*lh*_(*c, u*) denote the resulting attention weight from query-time bin *c* to key-time bin *u*, suppressing the session index. Attention width and excess width were defined by

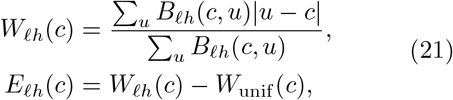

where *c* and *u* denote bin-center times in milliseconds, and *W*_unif_ (*c*) is the corresponding mean absolute lag under the geometry-only uniform-attention null. Subtracting this reference accounts for the dependence of expected lag on query position and available temporal context. Positive excess width indicates broader weighting than the null, and negative excess width indicates narrower weighting. Figure 5d displays standardized excess-width profiles.

#### Reference trajectory and head ranking

The example Integration–Split trajectory in Figure 1f was digitized by tracing the plotted curve, calibrating both axes to the panel tick marks, and resampling to the attention analysis’s 10-ms time bins. Let *R*(*c*) denote this reference. Correspondence for session *s*, layer *l*, and head *h* was quantified across query times as

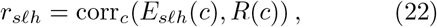

using Pearson correlation. Within each session and layer, heads were ranked in descending order of *r*_*slh*_, and maps and profiles were averaged across sessions at each rank. Head 1 therefore denotes the most strongly correlated head within a session and layer, not a fixed model head index. The *r* values above the maps are means of the session-level correlations for the corresponding ranks, not correlations computed after averaging profiles. Ranking was performed after training and did not modify model parameters or attention weights.

#### Control analyses

Matched, randomly initialized models with the same architecture and initialization seeds were analyzed using the same attention-extraction procedure. Potential dependence on spike density was assessed by comparing attention mass at each key time with the population peristimulus time histogram and recomputing widths after dividing attention mass by local spike density. To assess whether width reflected distance from one preferentially attended moment, each head’s excess-width profile was compared with |*t*^*\**^− *c*|, where *t*^*\**^ is its most-attended key time. Profiles were also compared with the independently estimated sliding-window decodability at matching query times (Figure 2a). Figure 5e additionally reports shuffled-label and shuffled-spike controls.

#### Single-head attention ablation

The contribution of learned temporal weighting was evaluated by intervening on one head at a time, following the general 20 use of head ablation to assess attention-head importance [36]. For query token *i* with valid-key set *K*_*i*_, its attention weights in the selected head were replaced by

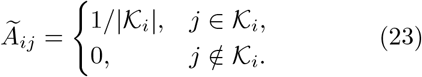

The value projections and other model parameters were retained. The manually implemented forward pass reproduced the unablated model’s output when no intervention was applied. For each session, layer, and head, the ablation effect was

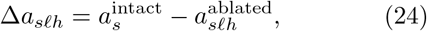

where both accuracies were evaluated on the same held-out trials. Positive Δ*a* indicates impaired decoding. Associations between *r*_*slh*_ and Δ*a*_*slh*_ were summarized using Spearman correlation and ordinary least-squares regression, pooled across the 88 head– session observations (11 sessions ×2 layers ×4 heads) and separately by layer (Figure 5f,g). The intervention tests the contribution of a head’s learned weighting as a whole; it does not selectively remove only the contraction-and-recovery component.

## Statistical analyses

Unless otherwise noted, optimal temporal scale, decodability, 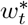, and Δ*R*^2^ are reported as mean ± SEM across the independent optimization or train/test repetitions, computed separately for each region, pseudo-session, and stimulus condition. Relationships between pooled time-resolved profiles were quantified with Pearson’s correlation using `scipy.stats.pearsonr`. These analyses yielded the *r* and *p* values reported directly in the Results.

To test whether a relationship was consistent across individual recordings and classification problems, Pearson’s *r* was also calculated separately for each pseudo-session/subtask case. The resulting distribution was tested against zero within each region using a one-sample *t*-test (`scipy.stats.ttest 1samp`); the Wilcoxon signed-rank test (`scipy.stats.wilcoxon`) was used as a non-parametric check. Significance markers in Figures 2f, 3l, 4g, and Supplementary Figure S1d,e denote the one-sample *t*-test. Tests were two-sided, the significance threshold was *α* = 0.05, and no multiple-comparison correction was applied to these region-level tests. Analyses were implemented in Python using NumPy, SciPy [41], and scikit-learn [42].

## Data and code availability

All neural data analyzed in this study are publicly available from the Allen Institute Brain Observatory: the Neuropixels Visual Coding dataset [30] and the Visual Behavior Neuropixels dataset [31]. Both were accessed programmatically through the AllenSDK and are subject to the Allen Institute’s Terms of Use. Code implementing the Integration–Split algorithm and the analyses described above will be available at `[repository URL]`.

## Supplementary Figures

**Supplementary Figure S1.**
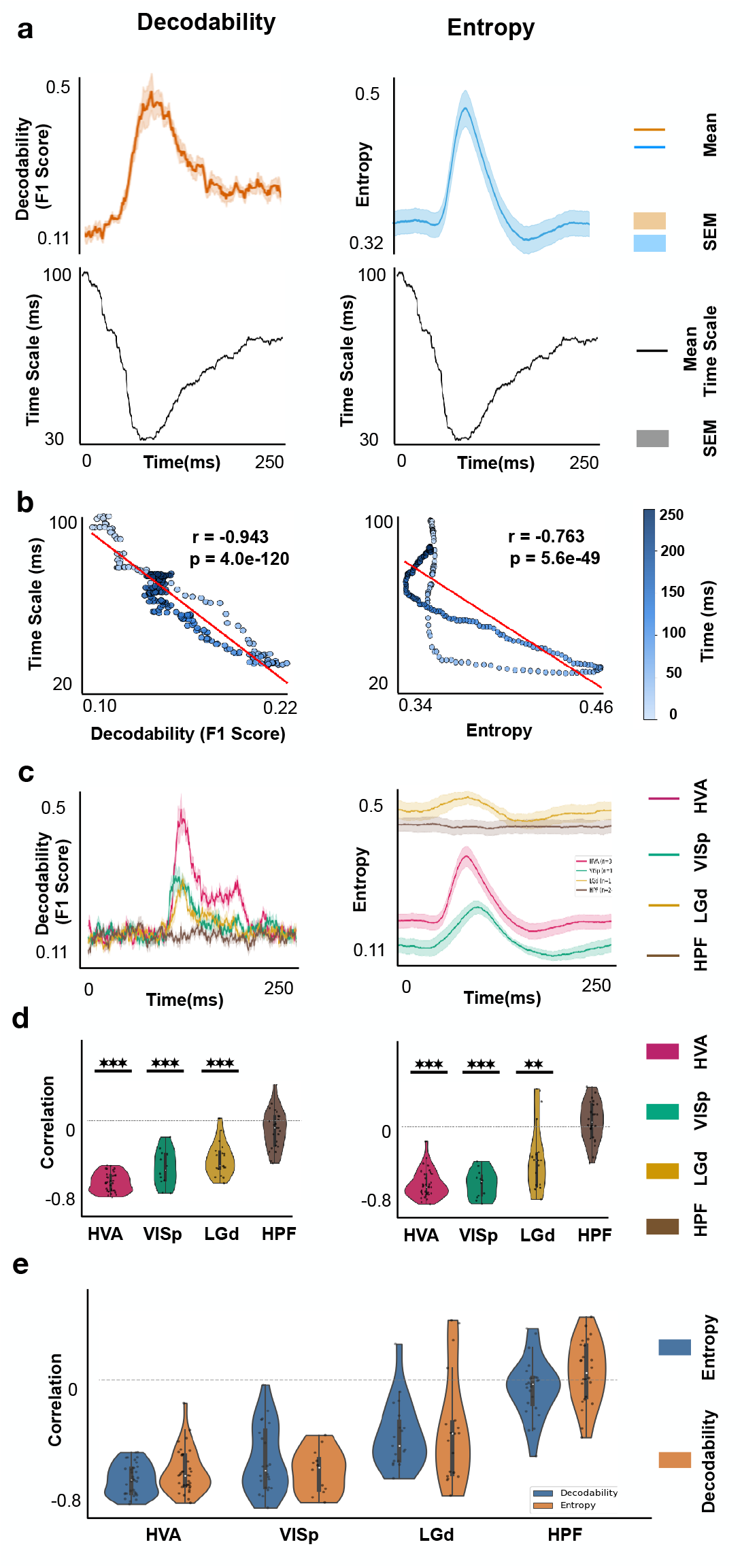
Response entropy tracks the optimal temporal scale but does not directly measure stimulus-specific information. Panels (a,b) show the Gabor paradigm in HVAs; panels (c–e) compare regions within the same paradigm. (a) Mean ± SEM decodability (*left*, orange) and response entropy (*right*, blue), with the corresponding Integration–Split optimal temporal scale below each profile. Both response entropy and decodability increase near the time at which the optimal temporal scale is finest. After the peak, entropy falls below its initial value even though decodability remains elevated, illustrating that response diversity and stimulus separability are not equivalent. (b) Optimal temporal scale plotted against decodability (*left* ; *r* = *−*0.943) and entropy (*right* ; *r* = *−*0.763) across time. Points are colored by time and red lines show least-squares fits. (c) Mean ± SEM decodability (*left* ) and entropy (*right* ) separated by region: HVAs (magenta), V1 (teal), LGd (gold), and HPF (brown). (d) Distributions of the per-case Pearson correlation between optimal temporal scale and decodability (*left* ) or entropy (*right* ) for each region. (e) Direct comparison of the decodability-based (orange) and entropy-based (blue) correlation distributions. No significant difference between the measures was detected within any region. In (d,e), points denote individual pseudo-session/classification cases; in (d), asterisks indicate a correlation distribution different from zero by a two-sided one-sample *t*-test (^*\**^*p <* 0.05, ^*\*\**^*p <* 0.01, ^*\*\*\**^*p <* 0.001; no multiple-comparison correction).

**Supplementary Figure S2:**
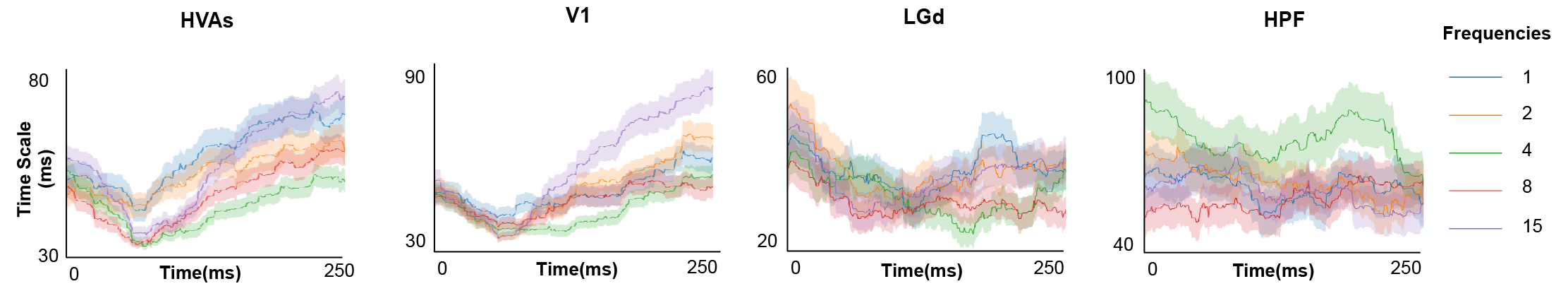
Optimal temporal-scale dynamics are not systematically ordered by drifting-grating frequency. Mean *±* SEM Integration–Split optimal temporal-scale profiles in HVAs, V1, LGd, and HPF, with drifting-grating trials analyzed separately at 1, 2, 4, 8, and 15 Hz. The visual regions retain the characteristic contraction-and-recovery profile across frequencies, whereas HPF remains comparatively weakly modulated. The relative ordering of the five profiles varies over time and across regions, indicating no consistent association between grating temporal frequency and the decoding-optimal temporal scale.

